# The RhoGAP myosin 9/HUM-7 integrates membrane signals to modulate Rho/RHO-1 during embryonic morphogenesis in *C. elegans*

**DOI:** 10.1101/325274

**Authors:** Andre G. Wallace, Hamidah Raduwan, John Carlet, Martha C. Soto

## Abstract

During embryonic morphogenesis, cells and tissues undergo dramatic movements under the control of F-actin regulators. Our studies of epidermal cell migrations in developing *C. elegans* embryos have identified multiple plasma membrane signals that regulate the Rac GTPase, thus regulating WAVE and Arp2/3 complexes, to promote branched F-actin formation and polarized enrichment. We describe here a pathway that acts in parallel to Rac to transduce membrane signals to control epidermal F-actin through the GTPase Rho. Rho contributes to epidermal migrations through effects on underlying neuroblasts. Here we identify signals to regulate Rho in the epidermis. HUM-7, the *C. elegans* homolog of human Myo9A and Myo9B, regulates F-actin dynamics during epidermal migrations, by controlling Rho. Genetics and biochemistry support that HUM-7 behaves as GAP for the Rho GTPase, so that loss of HUM-7 enhances Rho-dependent epidermal cell behaviors. We identify SAX-3/ROBO as an upstream signal that contributes to attenuated Rho activation through its regulation of HUM-7/Myo9. These studies identify a new role for Rho during epidermal cell migrations, and suggest that Rho activity is regulated by SAX-3/ROBO acting on the RhoGAP HUM-7.

## Introduction

Morphogenesis requires migrations of sheets of adherent cells. The migration and adhesion of the cells are under the control of GTPases. We have demonstrated that *C. elegans* embryos undergo morphogenetic movements that require the GTPase Rac1/CED-10 for epidermal enclosure, the sheet migration that leads to enclosure of the embryo by the six rows of epidermal cells that form at the dorsal surface. To coordinate these migrations, the GTPase Rac1/CED-10 activates the nucleation promoting factor WAVE/SCAR, which signals to the branched actin nucleator Arp2/3 to initiate actin polarization. Mutating any component in this pathway completely prevents the initiation of a sheet migration required for the enclosure of the embryo by epidermal cells during morphogenesis in *C. elegans* (Fig. 1A) (Patel et al., 2008). Our group was the first to show that three axonal guidance receptors, UNC-40/DCC, SAX-3/Robo and VAB-1/Eph, are required during embryonic morphogenesis to regulate the levels, localization and polarization of this actin nucleation pathway in epidermal cells (Bernadskaya et al., 2012). However, the molecules that connect the axonal receptors to the regulation of GTPase activity during this migration are not known.

**Fig. 1.**
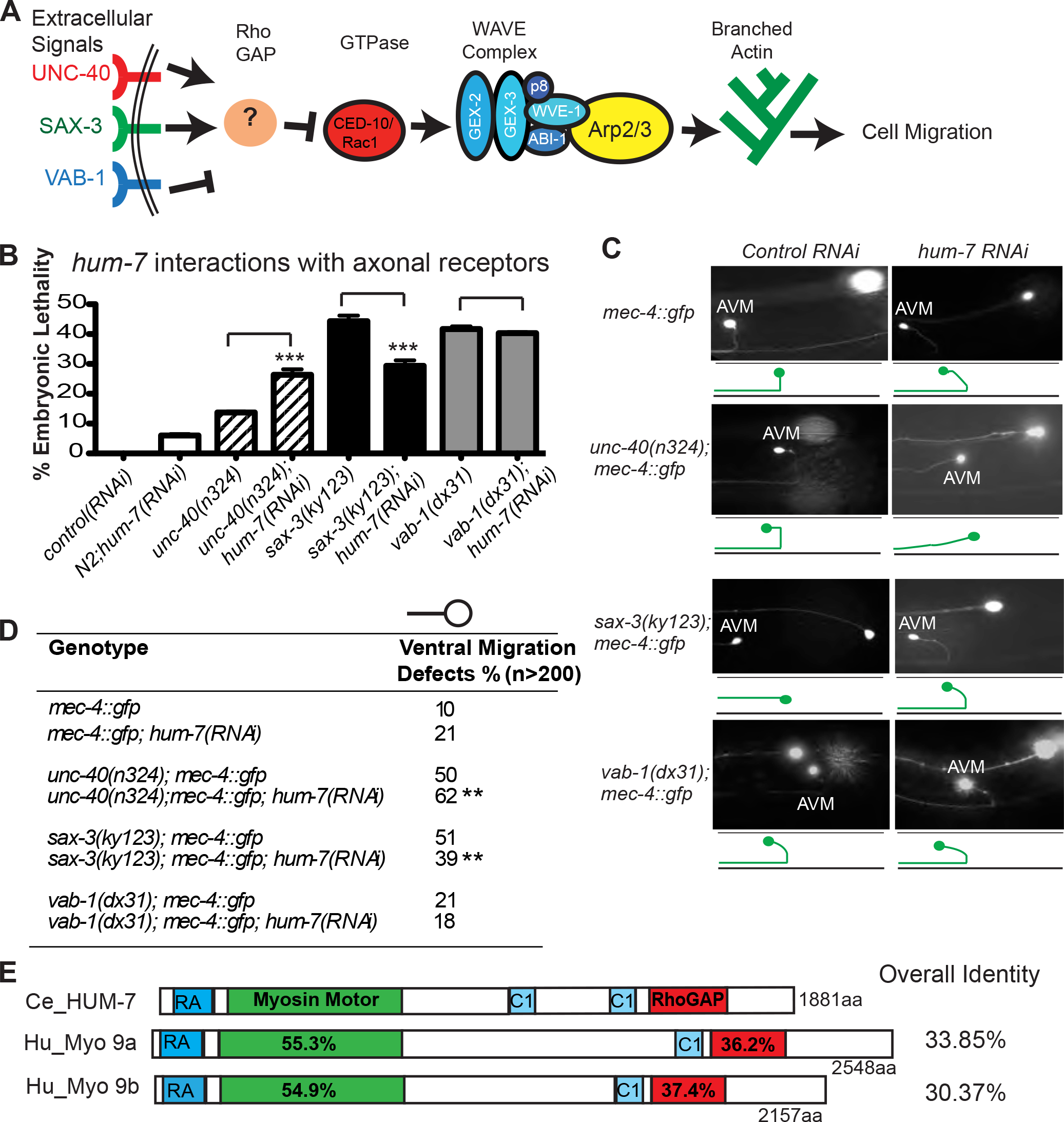
HUM-7/Myosin 9 interacts genetically with axonal guidance receptors and is required during embryonic morphogenesis. **A.** Axonal guidance receptors regulate the CED-10/Rac1 GTPase, which when active recruits the WAVE/SCAR complex to turn on Arp2/3, which promotes branched actin nucleation. Knock down of any WAVE/SCAR pathway component leads to full embryonic lethality, while the null mutations in upstream regulators result in partial embryonic lethality. To identify new regulators of morphogenesis, we screened for enhancers of *unc-40* embryonic lethality. **B.** The effects of loss of HUM-7 on embryonic lethality in axonal guidance receptor mutants. Genetic null mutants of three axonal guidance receptors (UNC-40, SAX-3 and VAB-1) were crossed into *dlg-1::gfp* to make scoring easier and treated with *hum-7* RNAi. Loss of HUM-7 enhanced *unc-40* lethality, suppressed *sax-3* lethality and did not alter lethality in *vab-1* mutants. The bar graph shows percent embryonic lethality. For this and all graphs shown in this study, error bars show SEM (Standard Error of the Mean). Statistical significance: *= p<.05, ** = p<0.001, *** = p<0.0001. For panel B statistical significance was determined by the One-way Anova test, followed by the Tukey test. Percentages are from three separate experiments. At least 500 embryos were analyzed in each experiment for each genotype. See also Table 1. **C.**Postembryonic effects of HUM-7 on regulators of the WAVE/SCAR pathway. The *mec-4::gfp (zdIs5)* transgene that is expressed in six mechanosensory neurons was used to show the effects of *hum-7* on axonal guidance receptors. In wild type animals the AVM axon migrates ventrally and then anteriorly, while mutants with defects in AVM axonal migration show initial migrations in other directions, including a complete loss of ventral migration (as is shown for the *sax-3(ky123)* animal). Micrographs show representative ventral migration patterns of the AVM axon, mimicked in the cartoons below (green cells). The black line in the cartoon represents the ventral nerve cord. The second neuron seen in some micrographs is the ALM, which is shown to orient the worm positioned anterior to the left. Control worms were fed L4440 empty vector RNAi. *hum-7* RNAi was monitored by counting the embryonic lethality in parallel. Worms for this experiment were grown at 20^°^C and only L4 stage larval worms were analyzed for neuronal assays. **D.** Table summarizing ventral migration AVM defects. Statistical significance was calculated in GraphPad Prism by a One-way ANOVA test followed by the Tukey test. Asterisks represent significance, *** = p<0.001. More than 200 neurons were analyzed for each genotype. **E.** *C. elegans* HUM-7 shares similar domains and sequence homology with human Myosin 9a and Myosin 9b. All three proteins possess a Ras-associated (RA) domain, which is embedded in the Ubitiquin (UBQ) domain of human Myosin-IXa, a myosin 9/IX domain and a RhoGAP domain. Both human Myosin 9 proteins have one phorbol ester/diacylglycerol-binding domain (C1) while HUM-7 has two. The percent sequence identity between *C. elegans* HUM-7 and the human Myosin 9 proteins are listed between domains (within labeled domains) and overall (on the right).

Recent studies by several groups suggest that besides Rac1, the GTPases RHO-1 and CDC-42 are also contributing to a morphogenetic process of ventral enclosure (Ouellette et al., 2016; Walck-Shannon et al., 2016; Wernike et al., 2016; Zilberman et al., 2017). However, the cytoskeletal consequences of altering RHO-1 or CDC-42 function during ventral enclosure are poorly understood. Alisa Piekny and colleagues have shown that myosin is required in the underlying neuroblasts to support epidermal cell migrations, suggesting important roles for Rho in neuroblasts during embryonic morphogenesis (Fotopoulos et al., 2013; Wernike et al., 2016). These studies raise the question of how Rho and myosin in the underlying neuroblasts influence the behavior or the migrating epidermals cells. A model has been proposed that myosin-based mechanical forces in neuroblasts influence constriction of the overlying epidermal cells. An alternative role for the neuroblasts is to promote a chemical cue that guides ventral enclosure, but the identity of this cue was not identified.

In *C. elegans* the GTPase RHO-1/Rho1 is needed throughout embryogenesis to promote cytokinesis, and also contributes to a step in *C. elegans* morphogenesis that occurs after the epidermis encloses the embryo, called embryonic elongation. During elongation the epidermis constricts circumferentially to squeeze the embryo into a tubular worm. Embryonic elongation also depends on regulators of Rho-1, and on Rho-1 targets including the Rho Kinase/LET-502 and non-muscle myosins (Chisholm and Hardin, 2005; Diogon et al., 2007; Fotopoulos et al., 2013; Gally et al., 2009; Martin et al., 2014; Piekny and Mains, 2003; Piekny et al., 2003; Quintin et al., 2008; Wissmann et al., 1997; Zhang et al., 2010). Elongation is driven by high levels of tension, mainly in the two rows of lateral seam cells. This tension is transmitted to the dorsal and ventral rows. Tension in the lateral cells is initiated by the Rho-binding kinase, LET-502 and its target, MLC-4/myosin regulatory light chain, which is highly expressed in the lateral cells. In contrast, the MEL-11/myosin phosphatase opposes this tension. While the lateral cells drive constriction, expression of the RhoGAP RGA-2 in the two rows of dorsal and ventral epidermal cells promotes decreased Rho activity and decreased tension (Diogon et al., 2007). This balance of forces acting on Rho-1 is required for proper elongation. It is therefore possible that changes in the balance of forces acting on Rho contributes to the cell movements of ventral enclosure.

Rho GTPases are regulated by GAPs, which enhance the hydrolysis of GTP to GDP, thus promoting dynamic turnover of GTPases from their active to inactive state. This turnover is important to modulate GTPase activity: studies suggest that specific levels of GTPases, and their subcellular enrichment, help promote correct signals that organize the actin cytoskeleton (Clay and Halloran, 2013). The ability of GAP proteins to turn over activated GTPases makes them attractive candidates to regulate GTPase function by controlling location and levels. However, there are many more GAP proteins than there are target GTPases, so removal of individual GAP proteins often gives subtle phenotypes that makes them harder to characterize. Removal of some GAPs that are known to result in strong, striking changes in the cytoskeleton in tissue culture cells results in mild or minimal effects when removed from developing organisms (Zaidel-Bar et al., 2010). Several GAPs are not essential for embryonic development, unless they are removed in the context of reduced apical junctions (Walck-Shannon et al., 2016; Zilberman et al., 2017).

The RhoGAP Myo9/Myosin IX is proposed to regulate multiple cellular functions including adhesion, cell migration and immune function. Mammals have only two of these myosins, Myo9a and Myo9b (Reviewed in Omelchenko 2012). Loss of function mutations or overexpression of these proteins has been linked to various cancers and immune defects. Some studies support a role for Myo9B in human intestinal diseases like inflammatory bowel disease and Crohn’s disease (Li et al., 2016; Prager et al., 2014; Wang et al., 2016). This conserved protein has a complex structure. Myo9 proteins have a C-terminal GAP domain and most studies support that this GAP domain can attenuate Rho activity. Myo9 proteins also have an N-terminal single headed processive myosin motor that can move the protein towards the plus or barbed ends of F-actin fibers (Chandhoke and Mooseker, 2012; Hanley et al., 2010; Hegan et al., 2016; Inoue et al., 2002; Liao et al., 2010; Omelchenko and Hall, 2012). The role of other conserved motifs, including RA (Ras associated) and C1 domains, is less well understood. The mouse knock out of Myo9A resulted in CNS defects (Abouhamed et al., 2009) and kidney defects (Thelen et al., 2015) that may also involve changes in protein trafficking, while the mouse Myo9b knockout resulted in altered morphology and motility of immune cells (Hanley et al., 2010) and impaired intestinal barrier function (Hegan et al., 2016). A pathway has been proposed in a human lung cancer model where SLIT/ROBO acts to inhibit Myo9b, which releases RhoA. However, the exact reason why this double inhibitory pathway is needed to regulate RhoA, and how misregulated RhoA promotes lung cancer is unclear (Kong et al., 2015). In metastatic prostate cancer cell lines Myo9b is elevated, while knockdown of Myo9b results in altered NMY2A (non muscle myosin), and defective migrations, further supporting a function in Rho attenuation (Makowska et al., 2015).

*C. elegans* has a single Myo9 homolog, HUM-7, which has been characterized extensively at the biochemical level, and was used to analyze how this single headed myosin motor is able to move processively towards the plus end of actin fibers using an insertion in the myosin head loop 2 (Liao et al., 2010). However, the *in vivo* role of HUM-7 has not been reported. *C. elegans* is thus an excellent *in vivo* system to probe the mechanisms behind the complex phenotypes seen for mutations in Myo9/HUM-7.

We sought novel regulators of morphogenesis and identified the Myo9/Myosin IX/HUM-7 protein. In this study we characterize the contribution of Myo9/HUM-7 to *C. elegans* morphogenesis. We investigate which GTPases are affected by this RhoGAP, and place *hum-7* in a genetic pathway known to regulate actomyosin contractility. In addition, we address how this proposed myosin IX motor affects F-actin levels, polarization and dynamics, and how it compares to known Rho pathway mutants, including myosin phosphatase, *mel-11*. We also investigate the contribution of Myo IX/HUM-7 to axonal guidance. Genetic epistasis studies place this GAP downstream of two receptors, SAX-3/ROBO and VAB-1/EphB, known to regulate F-actin during morphogenesis, to modulate Rho signaling. Targeted mutations and GFP tagging using CRISPR allow us to test genetic regulation and tissue expression of this RhoGAP.

## Results

Three axonal guidance receptors are upstream regulators of F-actin during morphogenesis through their effects on WAVE/SCAR (Fig. 1A) (Bernadskaya et al., 2012). To identify proteins that might connect the axonal guidance receptors to WAVE/SCAR regulation, we performed an RNAi screen to identify genes that enhance the low levels of embryonic lethality (14%) caused by loss of one of the axonal guidance receptors, UNC-40. We screened through a 2000-clone feeding RNAi library (Sieburth et al., 2005) which represents about 10% of all *C. elegans* genes and identified several enhancers of *unc-40* lethality using the putative null allele *unc-40(n324)*. RNAi depletion of HUM-7, an unconventional myosin heavy chain protein, increased *unc-40* embryonic lethality to 26%. *hum-7* was thus a candidate gene to act in parallel to *unc-40* during morphogenesis.

### HUM-7 functions in post-embryonic axonal guidance and genetically interacts with WAVE/SCAR axonal guidance regulators in embryos and larvae

To test if *hum-7* affects other signals that regulate the WAVE-1 complex similarly to how it affects *unc-40*, we fed *hum-7* RNAi to animals missing other axonal guidance receptors. One possible outcome might be enhancement of all the known signals that organize F-actin in developing embryonic epidermis. However, while loss of HUM-7 significantly increased the embryonic lethality caused by loss of UNC-40, it had no significant effect on the *vab-1(dx31)null* mutation, and it suppressed the embryonic lethality caused by the *sax-3(ky123)* null mutation from 44% to 29% (Fig. 1B). Therefore loss of *hum-7* had distinct effects on the three upstream signals that organize the cytoskeleton in developing embryos.

One human homolog of HUM-7, Myo9A, is mutated in patients with defects in the neuromuscular junction, leading to myasthenic syndrome (O’Connor et al., 2016). To test if HUM-7 had a post-embryonic developmental role, particularly in neurons, we characterized the loss of HUM-7 on six mechanosensory neurons, including the AVM neuron, that are easily visualized using the neuronal transgene *mec-4::gfp* (Yu et al., 2002). In a wild type background, the AVM neuron has a single round cell body and sends out one axonal projection that travels ventrally to the nerve cord and then anteriorly (Fig. 1C). RNAi depletion of HUM-7 in the *mec-4::gfp* transgenic strain led to doubling of AVM ventral migration defects. Therefore HUM-7 regulates development post-embryonically in the neurons known to be affected by the loss of the axonal guidance receptors UNC-40, SAX-3 and VAB-1.

To determine if *hum-7* and the axonal guidance receptor mutants interact genetically during postembryonic development, we analyzed the effects of *hum-7* RNAi on strains containing null alleles of the axonal guidance mutants and the *mec-4::gfp* transgene. The ventral migration defects caused by *unc-40(n324)* were enhanced from 50% to 62% by *hum-7* RNAi. In contrast, *sax-3(ky123)* ventral migration defects were suppressed from 51 % to 39% by *hum-7* RNAi. There were no significant differences in ventral migration defects in *vab-1(dx31)* mutants fed *hum-7* RNAi (Fig. 1C,D). Therefore, the post-embryonic genetic interactions paralleled the embryonic genetic interactions. These results suggest the effects described for HUM-7 function in the embryo are likely conserved postembryonically in other tissues, including migrating axons.

### HUM-7 functions in embryonic morphogenesis

If HUM-7 is an important regulator of morphogenesis, it should have a phenotype on its own. We depleted *hum-7* via RNAi in a wild type (N2) background, which resulted in 6.5% embryonic lethality. The *hum-7(ok3054)* mutation generated by the *C. elegans* Knockout Consortium includes a 653 bp deletion including the 3’ end of the myosin domain and two IQ domains (binding sites for EF-hand proteins including regulatory myosin light chains and calmodulin) (Bähler and Rhoads, 2002) (Fig. 2A). We out-crossed this strain three times, and detected 5.8% embryonic lethality. Since RNAi and the small deletion could be masking the true loss of function phenotype, we generated an almost full length deletion using CRISPR, *hum-7(pj62)*, and observed similar phenotypes and slightly higher embryonic lethality of 9.5%. Approximately half of the dead embryos displayed a phenotype known as Gex (gut on the <u>ex</u>terior), which occurs 100% of the time when any gene in the WAVE complex is knocked out. In a Gex embryo, the epidermal ventral migration step of morphogenesis fails and as a result internal organs like the pharynx and intestine (gut) are exposed and the embryo dies (Soto et al., 2002). Other dead embryos displayed a 2-fold arrest phenotype indicating a defect at the next morphogenesis step, elongation. All of these *hum-7* phenotypes were observed in both *hum-7* deletion mutants, and in the *hum-7* RNAi embryos, and in similar percentages (Fig. 2A-D, Table 1). This is similar to putative null mutations in upstream regulators of F-actin, like *unc-40*, which are only 14% embryonic lethal. Therefore, loss of *hum-7* alone results in partially penetrant and strong morphogenesis phenotypes.

**Fig. 2.**
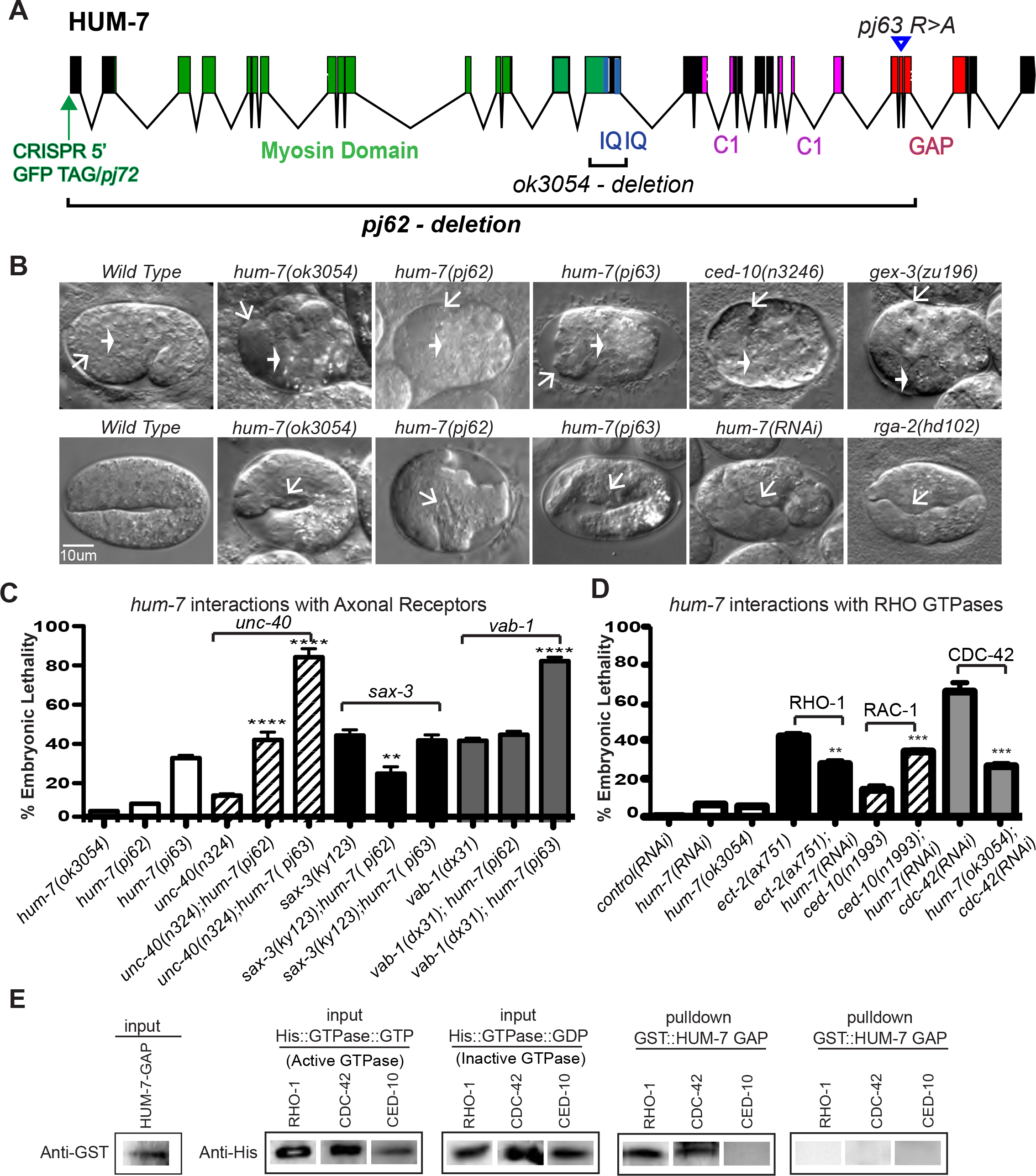
HUM-7/Myosin 9 interacts genetically and molecularly with Rho GTPases. **A.** Molecular model of the *C. elegans* Myosin 9 protein, HUM-7. The N-terminal Myosin, and C-terminal C1 and GAP domains are indicated. Genetic mutations from the C. elegans Consortium *(ok3054* deletion) or from our CRISPR studies *(pj62* deletion, *pj63* GAP mutant) are indicated, as well as the insertion site for an endogenous CRISPR N-terminal GFP tag, OX681 *hum-7(pj72 [gfp::hum-7]* (see Figure 3). **B.** Embryonic morphogenesis defects in *hum-7* mutants observed with DIC optics. Embryos are positioned with anterior to the left and dorsal up here and elsewhere unless otherwise noted. The thicker white arrows point to the anterior intestine while the thinner arrows point to the anterior pharynx. Some embryos missing *hum-7* arrest during ventral enclosure (top row), similar to embryos missing *ced-10/Rac1* or WAVE complex components like *gex-3*. Other *hum-7* embryos arrest at the 2fold stage (bottom row) similar to mutants that regulate Rho. All embryos shown in this study were mounted in Egg Salts (See Materials and Methods). **C.** Embryonic lethality percentages in genetic doubles of *hum-7* and axonal guidance mutants. CRISR was used to generate a large deletion, *pj62* and a R to A point mutation in the GAP domain, *pj63*, to study *hum-7* genetic interactions with null mutations in the axonal guidance receptors *unc-40, sax-3* and *vab-1*. At least 300 embryos were counted for each genotype. **D.** Embryonic lethality percentages in genetic and RNAi doubles of *hum-7* and GTPase mutants. We analyzed the three main *C. elegans* GTPases (RHO-1, CED-10 and CDC-42). We combined a hypomorphic allele of CED-10 *(ced-10(n1993))* with *hum-7* RNAi, RNAi of *cdc-42* with the *hum-7(ok3054)* mutant allele, and a hypomorphic allele of a RHO-1 guanine exchange factor *(ect-2(ax751))* with *hum-7* RNAi. The bar graph shows the percent embryonic lethality. At least three separate experiments were performed and more than 500 embryos per genotype was analyzed for each experiment. Error bars show SEM. Statistical significance: *** = p<0.001. See also Table 1. **E.** HUM-7 GAP domain binds to GTP-bound RHO-1 and GTP bound CDC-42 but not to GTP-bound CED-10. The GAP domain of HUM-7 was tagged with GST and used in a GST pull-down assay to test binding between three HIS-tagged GTPases (RHO-1, CDC-42 and CED-10) in both the active GTP-bound (Q63L, Q61L and Q61L) and inactive GDP-bound (T19N, T17N and T17N) states. The pull-down assays were performed with double the concentrations of the loading controls shown. The GST binding assay was done at least twice, using the same batch of GST-HUM-7-GAP for all of the pull-downs, and at least two sets of HIS-GTPase lysates. Details in the Materials & Methods.

**Table 1.**
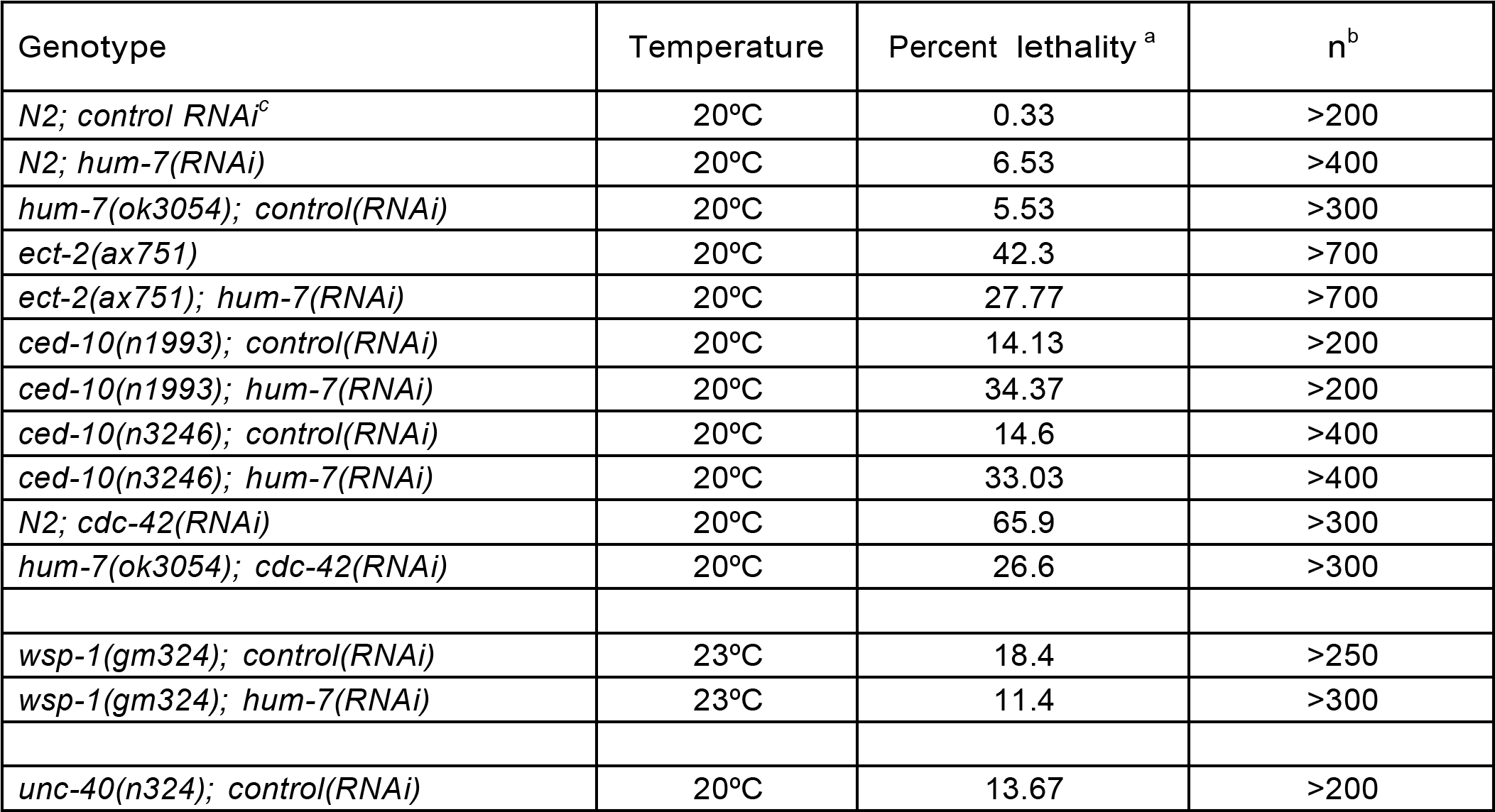
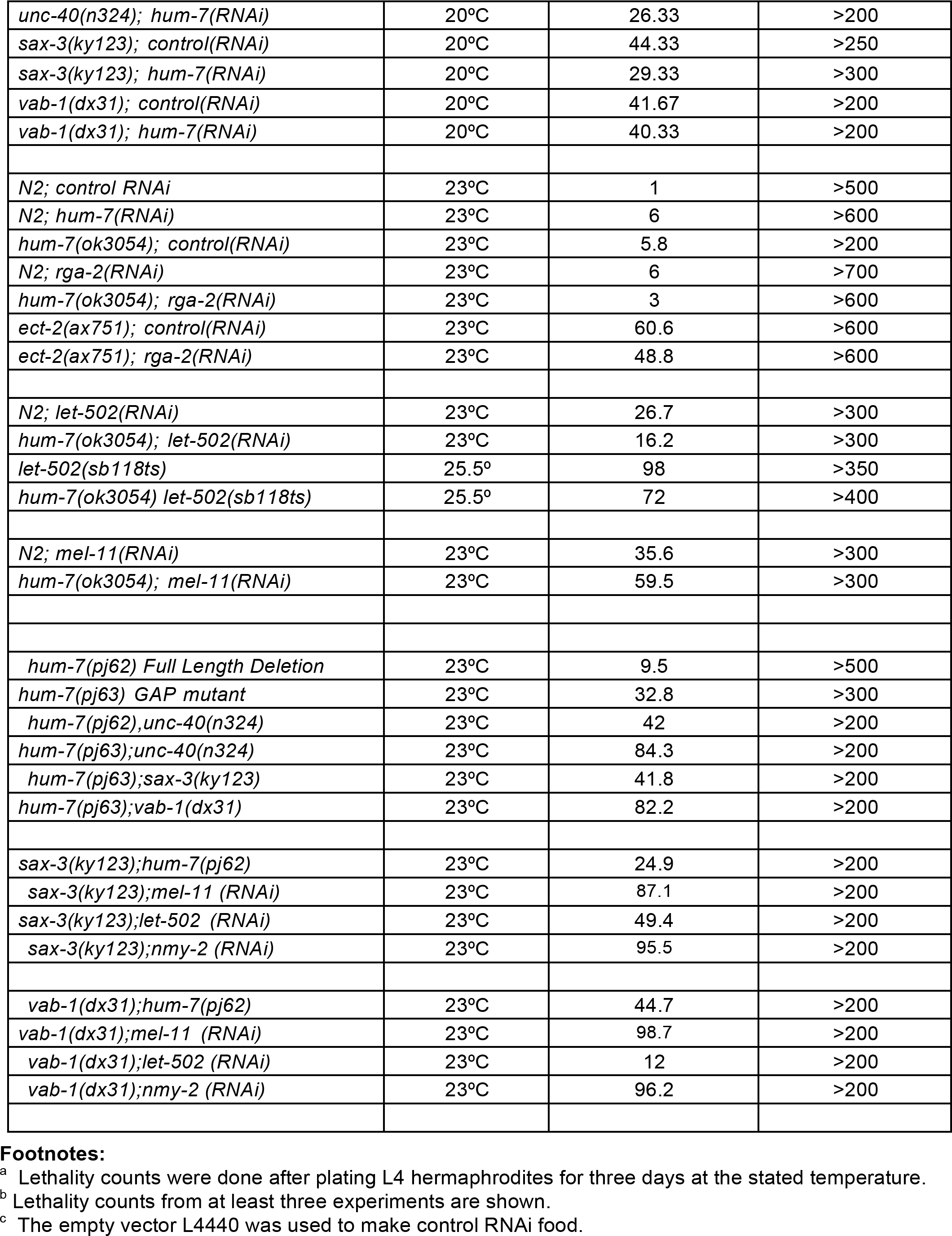
Embryonic lethality due to the loss of *hum-7* and *hum-7* double mutants. Related to Fig. 1B, 2C, D and Fig. 5A,B.

| Genotype | Temperature | Percent lethality <sup>a</sup> | n <sup>b</sup> |
| --- | --- | --- | --- |
| *N2; control RNAi*<sup>c</sup> | 20°C | 0.33 | >200 |
| *N2; hum-7(RNAi)* | 20°C | 6.53 | >400 |
| *hum-7(ok3054); control(RNAi)* | 20°C | 5.53 | >300 |
| *ect-2(ax751)* | 20°C | 42.3 | >700 |
| *ect-2(ax751); hum-7(RNAi)* | 20°C | 27.77 | >700 |
| *ced-10(n1993); control(RNAi)* | 20°C | 14.13 | >200 |
| *ced-10(n1993); hum-7(RNAi)* | 20°C | 34.37 | >200 |
| *ced-10(n3246); control(RNAi)* | 20°C | 14.6 | >400 |
| *ced-10(n3246); hum-7(RNAi)* | 20°C | 33.03 | >400 |
| *N2; cdc-42(RNAi)* | 20°C | 65.9 | >300 |
| *hum-7(ok3054); cdc-42(RNAi)* | 20°C | 26.6 | >300 |
| *wsp-1(gm324); control(RNAi)* | 23°C | 18.4 | >250 |
| *wsp-1(gm324); hum-7(RNAi)* | 23°C | 11.4 | >300 |
| *unc-40(n324); control(RNAi)* | 20°C | 13.67 | >200 |
| *unc-40(n324); hum-7(RNAi)* | 20°C | 26.33 | >200 |
| *sax-3(ky123); control(RNAi)* | 20°C | 44.33 | >250 |
| *sax-3(ky123); hum-7(RNAi)* | 20°C | 29.33 | >300 |
| *vab-1(dx31); control(RNAi)* | 20°C | 41.67 | >200 |
| *vab-1(dx31); hum-7(RNAi)* | 20°C | 40.33 | >200 |
| *N2; control RNAi* | 23°C | 1 | >500 |
| *N2; hum-7(RNAi)* | 23°C | 6 | >600 |
| *hum-7(ok3054); control(RNAi)* | 23°C | 5.8 | >200 |
| *N2; rga-2(RNAi)* | 23°C | 6 | >700 |
| *hum-7(ok3054); rga-2(RNAi)* | 23°C | 3 | >600 |
| *ect-2(ax751); control(RNAi)* | 23°C | 60.6 | >600 |
| *ect-2(ax751); rga-2(RNAi)* | 23°C | 48.8 | >600 |
| *N2; let-502(RNAi)* | 23°C | 26.7 | >300 |
| *hum-7(ok3054); let-502(RNAi)* | 23°C | 16.2 | >300 |
| *let-502(sb118ts)* | 25.5° | 98 | >350 |
| *hum-7(ok3054) let-502(sb118ts)* | 25.5° | 72 | >400 |
| *N2; mel-11(RNAi)* | 23°C | 35.6 | >300 |
| *hum-7(ok3054); mel-11(RNAi)* | 23°C | 59.5 | >300 |
| *hum-7(pj62) Full Length Deletion* | 23°C | 9.5 | >500 |
| *hum-7(pj63) GAP mutant* | 23°C | 32.8 | >300 |
| *hum-7(pj62);unc-40(n324)* | 23°C | 42 | >200 |
| *hum-7(pj63);unc-40(n324)* | 23°C | 84.3 | >200 |
| *hum-7(pj63);sax-3(ky123)* | 23°C | 41.8 | >200 |
| *hum-7(pj63);vab-1(dx31)* | 23°C | 82.2 | >200 |
| *sax-3(ky123);hum-7(pj62)* | 23°C | 24.9 | >200 |
| *sax-3(ky123);mel-11 (RNAi)* | 23°C | 87.1 | >200 |
| *sax-3(ky123);let-502 (RNAi)* | 23°C | 49.4 | >200 |
| *sax-3(ky123);nmy-2 (RNAi)* | 23°C | 95.5 | >200 |
| *vab-1(dx31);hum-7(pj62)* | 23°C | 44.7 | >200 |
| *vab-1(dx31);mel-11 (RNAi)* | 23°C | 98.7 | >200 |
| *vab-1(dx31);let-502 (RNAi)* | 23°C | 12 | >200 |
| *vab-1(dx31);nmy-2 (RNAi)* | 23°C | 96.2 | >200 |

<sup>a</sup> Lethality counts were done after plating L4 hermaphrodites for three days at the stated temperature.
<sup>b</sup> Lethality counts from at least three experiments are shown.
<sup>c</sup> The empty vector L4440 was used to make control RNAi food.

The *hum-7* CRISPR alleles and Consortium deletion were used to further probe *hum-7* genetic interactions with the axonal guidance genes. Testing the genetic interactions of *hum-7* using the two deletions, *ok3054* or *pj62*, gave similar results in combination with null mutations in *unc-40* (increased lethality), *sax-3* (suppressed lethality) or *vab-1* (no change) as we saw with *hum-7* RNAi (Figure 2C). This suggests RNAi results in strong loss of *hum-7* function. To test the importance of the GAP domain, we used CRISPR to generate a point mutation in the GAP of *hum-7, pj63*, and found different effects. This Arginine to Alanine (R to A) mutation is predicted to perturb the GTPase activating function of RhoGAPs, resulted in increased lethality compared to the deletions (30%), and more than additive lethality in combination with *unc-40* and *vab-1*. Unlike the deletion alleles, the *pj63* GAP mutant could not rescue *sax-3* embryonic lethality (Figure 2C). This result suggests the *hum-7* GAP mutant behaves as a dominant negative (worse than the large deletion) and further supports an interaction between *hum-7* and *sax-3*.

### HUM-7 functions as a GAP for Rho-1 and Cdc-42, but not for CED-10/Rac1

HUM-7 has strong sequence homology with myosin IX/Myo9 proteins from a variety of organisms, and includes the typical N-terminal myosin and C-terminal Rho GAP domains (Abouhamed et al., 2009; Chandhoke and Mooseker, 2012; Hanley et al., 2010; Omelchenko and Hall, 2012). HUM-7 has greater than 30% overall protein identity with human Myosin 9a and Myosin 9b (Fig. 1E). While a *C. elegans* GAP that regulates the GTPase CED-10 during cell corpse engulfment has been identified (Neukomm et al., 2011; Neukomm et al., 2014), this Rho GAP, SRGP-1, is completely dispensable for embryonic viability, and has limited morphogenesis phenotypes as a single mutant (Zaidel-Bar et al., 2010). We therefore tested if HUM-7 functions as a GAP for one or more *C. elegans* GTPases.

#### Genetic evidence that HUM-7 functions as a GAP for Rho-1 and Cdc-42, but not for CED-10/Rac1

We hypothesized that if HUM-7 acts like a GAP for CED-10/Rac1, then loss of *hum-7* would rescue partial loss of CED-10/Rac1. However, while the *ced-10(n1993)* hypomorphic mutation resulted in 14% embryonic lethality, depletion of *hum-7* via RNAi in the *ced-10(n1993)* hypomorphic allele increased embryonic lethality to 34% (Table 1, Fig. 2D). Experiments with a second hypomorphic allele, *ced-10(n3246)* similarly resulted in increased,not decreased embryonic lethality (Table 1). Thus *hum-7* and *ced-10* appear to function in parallel pathways.

To test if HUM-7 can function as a GAP for RHO-1/Rho, we used a temperature sensitive allele of the known Rho GEF, *ect-2(ax751)*. This allele has been well characterized to partially knock down Rho signaling (Zonies et al., 2010). At 20°C, *ect-2(ax751)*/*RHO-GEF* resulted in 42% embryonic lethality on control RNA, that dropped to 28% lethality in animals also depleted of *hum-7* via RNAi. Similarly, *ect-2* RNAi leads to almost 100% dead embryos but this drops to 84% dead embryos if RNAi is fed to *hum-7* mutants (Fig. 2D, Table 1). Further, all of the *ect-2* phenotypes, including those affecting the early embryo, were suppressed by the loss of *hum-7*.

To test if HUM-7 can function as a GAP for CDC-42, we used RNAi depletion of *cdc-42*, which in wild type animals resulted in embryonic lethality above 60%. In contrast, depletion of *cdc-42* via RNAi in the putative null strain *hum-7(ok3054)* resulted in embryonic lethality of only 27% (Table 1, Fig. 2D). The fact that loss of *hum-7* was able to suppress embryonic defects caused by partial loss of *cdc-42* or *rho-1* (using *ect-2(ax751)*) suggested that *hum-7* acts in a pathway with *rho-1* and *cdc-42*. Since there are no other reports of a Myosin 9 protein regulating *cdc-42*, we also tested if loss of *hum-7* could suppress the lethality of a known *cdc-42* effector, *wsp-1*. The deletion mutation *wsp-1(gm324)* results in 18.4% embryonic lethality that drops to 11.4% when these animals are depleted of *hum-7* via RNAi (Table 1).

*Molecular evidence that HUM-7 functions as a GAP for Rho-1 and Cdc-42, but not for CED-10/Rac1*. To test if the HUM-7 GAP domain can bind to the activated form of specific GTPases we performed GST pull-down assays. Purified His-tagged GTPases in active (GTP loaded) and inactive (GDP-loaded) form were individually tested for binding to the GAP domain of HUM-7. The GAP domain of HUM-7 bound strongly to GTP-loaded RHO-1 and CDC-42 and failed to bind GTP-loaded CED-10. No binding was observed between HUM-7 GAP and any of the GDP-loaded constructs tested for all three GTPases (Fig. 2E). Therefore biochemical studies suggested that HUM-7 could function as a GAP for RHO-1 and CDC-42 but not for CED-10. In short, HUM-7 behaves functionally, as well as molecularly, as a GAP for RHO-1 and CDC-42, but not CED-10.

### HUM-7 is expressed broadly with enhanced expression in muscles

If HUM-7 is important for embryonic development, we expect it to be expressed in embryos. We therefore created an N-terminally endogenously tagged allele of *hum-7, pj72, via CRISPR*. CRISPR tagged *gfp::hum-7* is broadly expressed in embryos at all stages, with robust, enhanced expression in muscle, including the developing body wall muscles (Fig. 3A). Staining *gfp::hum-7* expressing embryos with monoclonal antibody 5.6 (Miller et al., 1983) which recognizes body wall muscles, confirmed the muscle enrichment in embryos and larvae (Figure 3A, indicated by enlarged boxed region).

**Fig. 3.**
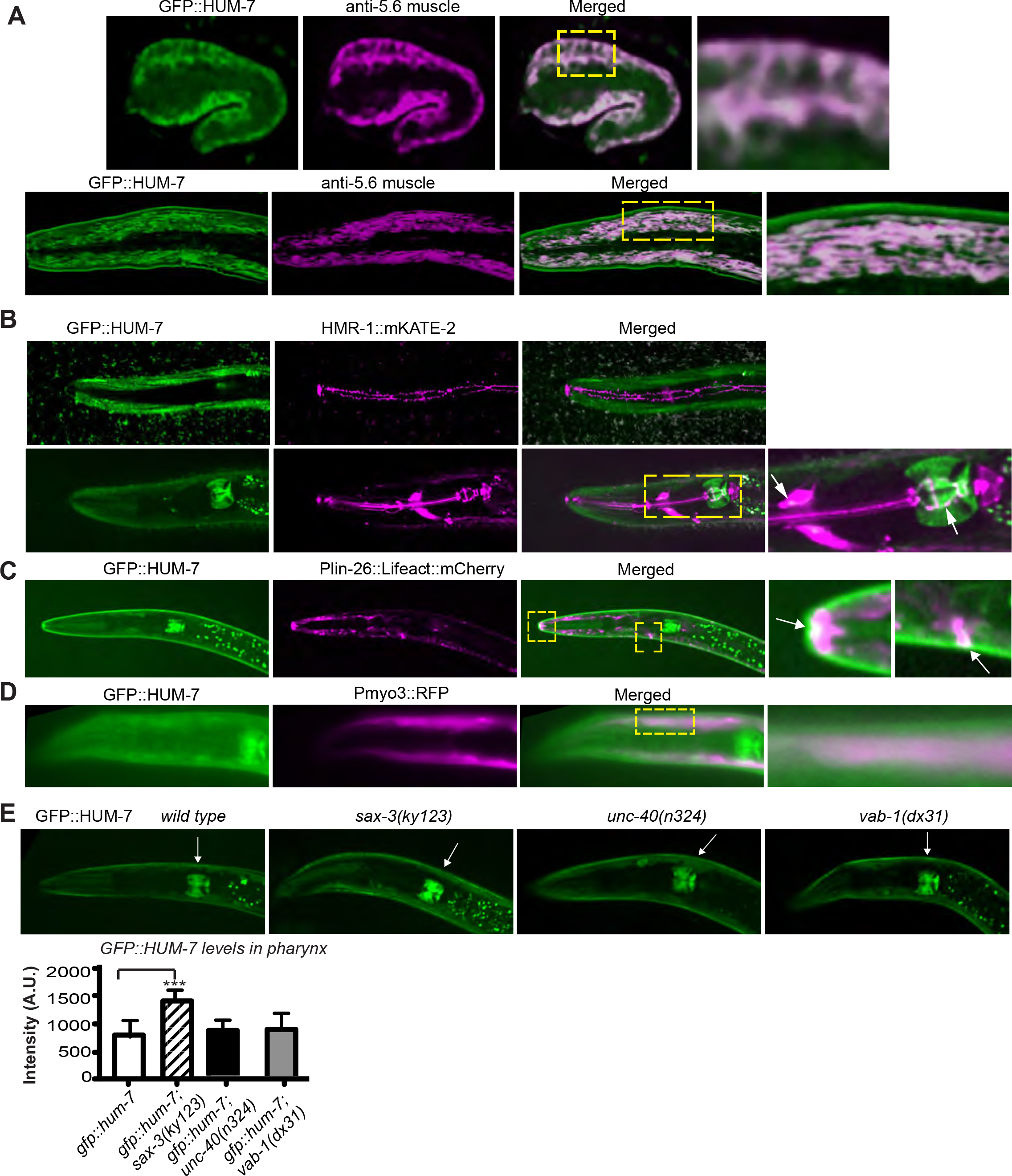
HUM-7 is broadly expressed, enriched in muscle, and regulated by SAX-3/ROBO. **A**. GFP::HUM-7 shows broad, diffuse expression in all tissues. However, particularly high expression is seen in the developing muscles, as shown by double labeling with anti-muscle antibody 5.6 (Miller et al., 1983). In larvae, the expression mirrors the striated appearance of the muscle fibers. Adult GFP::HUM-7 expression was compared with (**B**)HMR-1::mKATE2, (**C**) Plin26::Lifeact::mCherry and (**D**) Pmyo-3::RFP to compare tissue distribution. HMR-1/E-Cdh is known to be enriched at apical regions of epithelia and in the nerve ring. Pmyo-3::RFP is enriched in body wall muscles. Plin-26::Lifeact is enriched in the epidermis. **E**. The axonal guidance protein SAX-3/ROBO regulates levels of GFP::HUM-7. Arrows point to the posterior bulb of the pharynx. Maximum intensity at the pharynx was compared and graphed.

The *gfp::hum-7* CRISPR strain was crossed to other strains to analyze expression in other tissues. A strain carrying *gfp::hum-7* and *hmr-1::mKate2* (Marston et al., 2016) allowed us to compare expression in the central nervous system, pharynx and epidermis (Fig. 3B). In the posterior pharyngeal bulb *gfp::hum-7* is expressed all throughout, including the pm6 and pm7 cells that help the grinder contract during feeding [Worm Atlas], while *hmr-1::mKate2* is enriched only in apical regions, as expected. *hmr-1::mKate2* is expressed all through the nerve ring axons, while *gfp::hum-7* is expressed in adjacent cells, perhaps glia. Viewing this same strain on the surface shows that while *hmr-1::mKate2* is highly expressed in the seam cells, *gfp::hum-7* is instead enriched in the muscle cells, and shows no obvious overlap in the seam cells (Fig. 3B). A strain carrying *gfp::hum-7* and *Plin-26::Lifeact::mCherry* (Havrylenko et al., 2015), which expresses Lifeact under an epidermal promoter, similarly shows no overlap in the epidermal seam cells. An interesting overlap is around what appears to be the pore cell that connects the excretory cell to the outside (Figure 3C). This may be epithelial tissue, as interfacial epithelial cells are located at the junctions where the epidermis meets other types of tissues (Worm Atlas). A strain expressing RFP under control of the myosin heavy chain promoter, *Pmyo-3::rfp* (Viveiros et al., 2011), shares expression with *gfp::hum-7* in body wall muscles (Fig. 3D). Therefore *gfp::hum-7* appears to have broad expression, with enhanced expression in several types of muscle tissue including regions of the pharynx, and in body wall muscles. Expression is not overall enhanced in neurons, but may include support cells for neurons. Expression does not appear enhanced in epidermal cells, although there is precedent for epidermal signal to appear striped due to impingement from muscle or pharynx (Worm Atlas). Antibody staining with antibodies specific to body wall muscle support that at least some of the striped signals are in muscle (Figure 3A).

### sax-3 regulates levels of gfp::hum-7

*hum-7* genetic interactions with the axonal guidance receptors led us to test if any of them might be upstream regulators of *gfp::hum-7*. Loss of *unc-40* or *vab-1* had limited effects on *gfp::hum-7*, but loss of *sax-3* resulted in significantly elevated levels of *gfp::hum-7* in all tissues. Measuring the effect on the posterior bulb shows an increase of 100% (Fig. 3E). Therefore, the genetic and molecular interactions suggest *sax-3* is upstream of *hum-7* and that SAX-3 signaling results in lower HUM-7 expression.

The phenotypes, genetic interactions and biochemical studies suggested *hum-7* may function in the regulation of Rho during morphogenesis. To address how this proposed RHO-1 GAP, HUM-7, contributes to morphogenesis, we analyzed its contribution an essential event in embryonic development: ventral enclosure, which is driven by epidermal cell migrations, and is mainly thought to be under Rac1/CED-10 control.

### HUM-7 affects F-actin dynamics during the initial cell migrations required for epiboly

Some of the embryonic phenotypes resulting from loss of HUM-7 resemble the Gex phenotype seen when members of the WAVE/SCAR complex are mutated (Fig. 1A, 2B). Loss of WAVE/SCAR components during epidermal cell migration leads to defects in F-actin levels, ventral F-actin enrichment and actin dynamics in the migrating epidermis (Bernadskaya et al., 2012; Patel et al., 2008). We therefore tested HUM-7 effects on the actin cytoskeleton using two epidermal F-actin strains, *plin-26::vab-10 ABD(actin binding domain)::gfp (mcIs51)* transgene (Gally et al., 2009) and *plin26::Lifeact::mCherry* (Havrylenko et al., 2015). These transgenic strains, which express an actin binding protein (*vab-10* ABD or Lifeact) under the *lin-26* epidermal promoter, enabled us to perform live imaging of actin levels, ventral enrichment and dynamics in the epidermis of embryos during morphogenesis. In wild-type embryos during the early stages of morphogenesis (250 minutes after the first cell division), F-actin becomes enriched at the ventral edge of the leading cells of the epidermis. As morphogenesis proceeds F-actin levels increase further and continue to be ventrally enriched. When WAVE components are depleted via genetic mutation or RNAi overall F-actin levels drop (Patel et al., 2008). In *hum-7* mutants or *hum-7(RNAi)* embryos the overall levels of F-actin are higher than in wild type using the *plin-26::vab-10 ABD strain*, or the *plin26::Lifeact::mCherry* strain (Figure 4A-E). We conclude loss of *hum-7* increases epidermal F-actin levels.

**Fig. 4.**
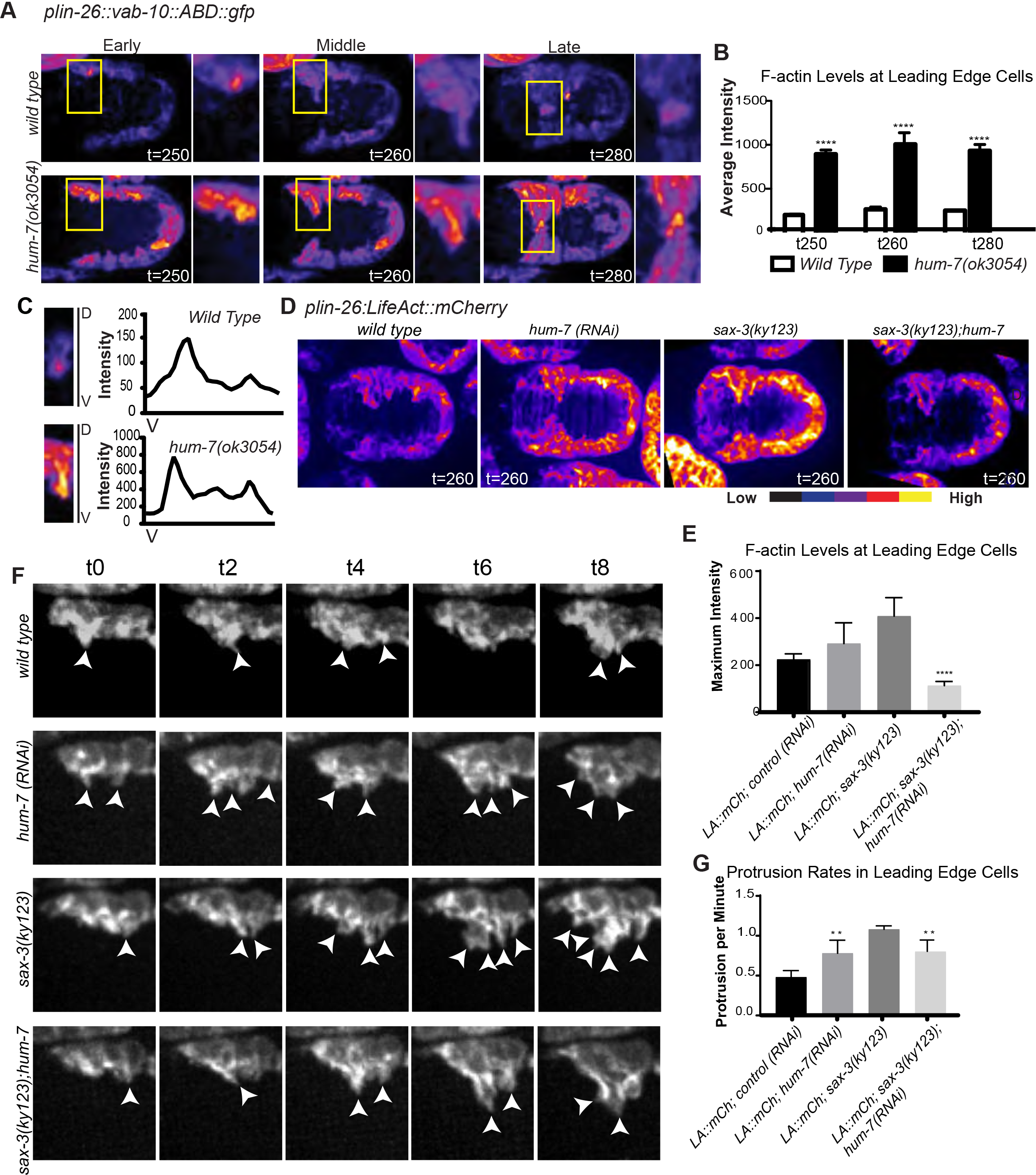
HUM-7 affects F-actin levels and dynamics in migrating epidermis during embryonic morphogenesis. F-actin levels in embryos during epidermal enclosure as shown by *plin-26::vab-10(ABD)::gfp<* (Gally et al., 2009),for panels A-C and *plin-26::Lifeact::mCherry* (Havrylenko et al., 2015) for panels D-G. **A,B**. Epidermal F-actin levels increase in *hum-7* mutants, *as shown by plin-26::vab-10(ABD)::gfp*. Embryos were imaged at two-minute intervals beginning at 240 minutes after first cleavage. Images shown here represent embryos at the early (250 min), middle (260 min) and late (280 min) stages of the ventral migration of epidermal enclosure. The embryos are oriented with anterior to the left and ventral up. The yellow boxes highlight the leading cells that lead the migration, and the average signal was measured. Embryos were pseudocolored using the “Fire” function in Image J, and intensity is shown from low (blue) to high (yellow). See also S1 Movie. **C**. Ventral enrichment of F-actin in migrating epidermal cells is maintained in *hum-7(ok3054)* embryos. Close-ups of representative epidermal leading cells with the ventral (V) edge at the bottom and the dorsal (D)region at top are shown. Intensity of F-actin in the leading cells during protrusion initiation was analyzed with a ventral to dorsal line through the leading cells using the Plot Profile tool in ImageJ. Note that the two graphs use different y-axis scales due to higher F-actin levels in *hum-7(ok3054)*. At least 10 embryos were analyzed per genotype. See also S2 Movie. **D**. *plin-26::Lifeact::mCherry* transgenic line (Havrylenko et al., 2015), in control animals and embryos depleted of *hum-7, sax-3* or both. **E**. The maximum F-actin intensity was measured in the ventral leading edge cells at 260 minutes after first cleavage. Analysis in IMAGEJ, using the “line tool”, drawn across the brightest region of the leading cells recorded the highest fluorescence intensity. At least 8 embryos were measured per genotype. **F, G**. Effects of *hum-7* and *sax-3* on the rate of dynamic actin protrusions. Using the same 4D movies shown in Panel D, the rate of F-actin protrusions was calculated in the two leading cells, which form frequent protrusions and retractions. The images show the two leading cells on one side of the embryo. Arrows mark protrusions and asterisks mark retractions. Beginning at 250 minutes after first cleavage, the number of protrusions occurring within an 8-minute period was recorded. Error bars show SEM and asterisks represent statistical significance, determined by One-way ANOVA followed by Tukey test *** = p<0.001. See also S2 Movie.

In WAVE mutants F-actin fails to enrich ventrally in the migrating cells and therefore fails to drive ventral enclosure (Bernadskaya et al., 2012; Patel et al., 2008). However, in *hum-7* mutants, F-actin is correctly enriched ventrally in the leading cells (Fig. 4C 10+ embryos per genotype). Therefore, *hum-7* does not appear to regulate ventral enrichment.

We next compared actin dynamics in *hum-7* mutants, crossed into both F-actin reporters. Previous work using the *plin-26::vab-10 ABD* transgene showed that wild type ventral epidermal cells undergo frequent protrusions and retractions at the edge of the leading cells, while protrusions and retractions in WAVE/SCAR mutants occurred less frequently (Bernadskaya et al., 2012; Patel et al., 2008). To assess the effects of *hum-7* on the actin protrusion dynamics, we imaged F-actin every two minutes from 240 minutes to 400 minutes after first cleavage. Wild-type embryos exhibited dynamic protrusions at the edge of the leading cells, with an average of 0.45 protrusions per minute (n= at least 10 embryos per phenotype). However in *hum-7* mutants, protrusions were more frequent, with an average of 0.8 protrusions per minute (S1 Movie, S2 Movie). This result was also seen with the second strain, *plin26::Lifeact::mCherry*, which showed an increase from 0.5 to 0.8 per minute (Fig. 4F,G). Therefore, the rate of actin protrusions is greater in *hum-7* mutants than in wild type, the opposite of the defect seen in WAVE mutants.

F-actin enrichment and dynamics were affected in *sax-3* mutants, and this depended on *hum-7*. Previous work using *plin-26::vab-10 ABD* showed that *sax-3(ky123)* embryos have relatively wild type levels of F-actin at the beginning of ventral enclosure, that becomes more disrupted as enclosure proceeds. The enclosure process was also delayed (Bernadskaya et al., 2012). When *sax-3(ky123)* was crossed into *plin26::Lifeact::mCherry* we observed elevated F-actin as the embryos enclosed. The double mutant with *hum-7* resulted in lower levels (Fig. 4D,E). Protrusions per minute were elevated in *sax-3* mutants, and this number also dropped in the double with *hum-7* (Fig. 4F,G). In both cases, loss of *hum-7* suppressed a *sax-3* F-actin phenotype in the epidermis.

### HUM-7 regulates timing of epidermal morphogenesis

Mutations in WAVE/SCAR, or in upstream regulators of WAVE/SCAR, including UNC-40/DCC/Netrin Receptor, SAX-3/Robo Receptor and VAB-1/Ephrin Receptor, lead to delayed migrations (Bernadskaya et al., 2012). We compared the timing of enclosure in *hum-7* mutants and found that while a low percent of embryos arrest partway through enclosure, embryos on average are not delayed, and some can enclose more quickly than wild type. For example, in one set of experiments 1/7 embryos arrested, while 2/7 embryos enclosed significantly faster than wild type. In Movies S1 and S2 the *hum-7(ok3054)* embryo in the center achieves ventral enclosure faster than wild type while the *hum-7(ok3054)* embryo on the right arrests during enclosure. We saw similar results, faster migrations, in the other *hum-7* mutants, *pj62* (20% faster) and *pj63* (10% faster) when compared to wild type embryos and timed from 240 minutes after first cleavage to ventral enclosure, by 300 min.). Together, these results demonstrate a role for HUM-7 in the regulation of F-actin dynamics that contribute to the correct initiation of epidermal cell migrations during epidermal enclosure, and correct timing of morphogenetic events.

### HUM-7 functions as a GAP upstream of RHO-1 to regulate embryonic morphogenesis

The role of the GTPases Rho/RHO-1 and Cdc42/CDC-42 in the ventral enclosure step of morphogenesis is only beginning to be described (Fotopoulos et al., 2013). In contrast, the role of RHO-1 in a later morphogenetic step, epidermal elongation, is exceedingly well studied (Diogon et al., 2007; Gally et al., 2009; Martin et al., 2014; Piekny et al., 2003; Piekny et al., 2000). Since *hum-7* behaved genetically and molecularly like a candidate GAP for RHO-1 (and CDC-42), we investigated if *hum-7* mutants have defects in Rho-dependent processes, like elongation. As shown in Fig. 2B, half of dying *hum-7* embryos show defects in elongation, including swelling of the anterior region. This defect is shown by known RHO-1 pathway mutants, including *rga-2*, a RHO-1 GAP that acts during elongation but not during ventral enclosure (Diogon et al., 2007) (thin arrows, Fig. 2B,Table 1).

If *hum-7* is a GAP for RHO-1, loss of HUM-7 might suppress mutations that decrease Rho-dependent contractility, like *let-502/RHO-1* Kinase, and enhance mutations that increase Rho-dependent contractility, like mel-11/Myosin Phosphatase. These genetic interactions have been demonstrated for the RHO GAP *rga-2* (Diogon et al., 2007). The *let-502 ts* allele, *sb118ts*, resulted in 75% or 98% lethality at 25°C or 25.5°C, respectively. However this lethality dropped to 60% or 72%, respectively, when we crossed in *hum-7(ok3054)* (Table 1 and Fig. 5A). In a wild-type background, *mel-11* RNAi resulted in 36% lethality and this lethality was enhanced to 60% in a *hum-7* background (Table 1, Fig. 5A). These results support that *hum-7* may be a GAP for RHO-1.

**Fig. 5.**
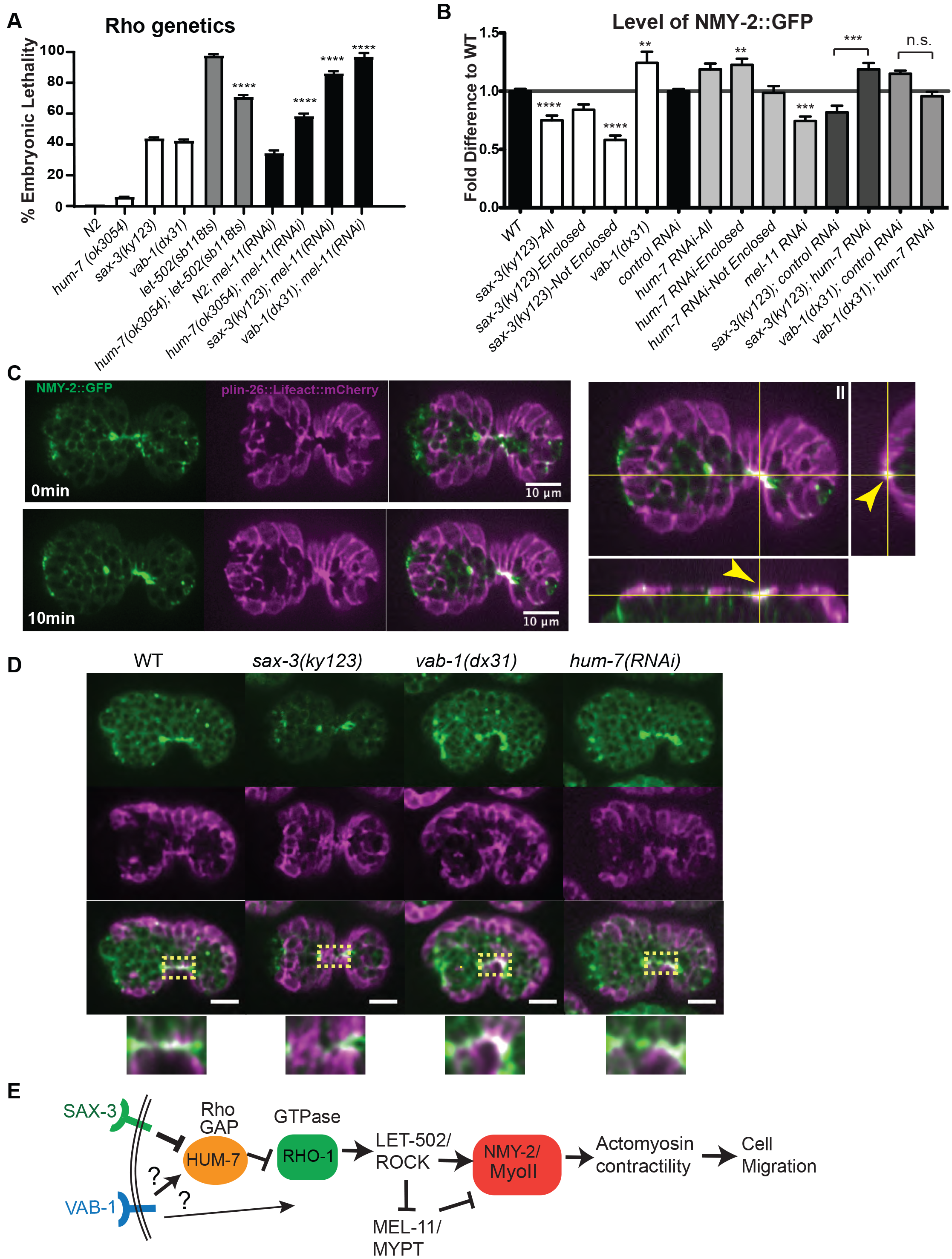
SAX-3/HUM-7/RHO-1 in epidermal cells regulate non-muscle myosin, NMY-2, to promote ventral enclosure. **A**. Genetic interactions of *hum-7, sax-3, vab-1* and known Rho pathway mutants Rho Kinase/*let-502* and myosin phosphatase/*mel-11*. Embryonic lethality was scored in single and double mutants using genetic mutants and RNAi depletion. Experiments shown were done at 23 C except for the *let-502(sb118ts)* experiments done at 25.5 C. **B**. Quantitation of levels of *nmy-2::gfp* in the ventral pocket cells at approximately 320 minutes after first cleavage, a time when control embryos have met at the ventral midline. See D for details. **C**. The CRISPR endogenously tagged *nmy-2::gfp* was crossed into *Plin-26::Lifeact::mCherry* to examine the expression of *nmy-2:gfp* during ventral enclosure. Colocalization in the epidermis is shown. The same embryo is shown 10 minutes apart, to show the colocalizing signal meets at the midline. Right panels show a 3D projection to better show the signals on the surface of embryos. **D**. *nmy-2:gfp; Plin-26::Lifeact::mCherry* was used to measure myosin enrichment in the ventral epidermal cells during different stages of ventral enclosure. Embryos during pocket cell alignment at the ventral midline. For embryos that do not enclose, ventral cells were measured at a similar time point. Yellow boxes indicate the region that was measured in B. **E**. Proposed model for how SAX-3, VAB-1 and HUM-7 regulate the RHO-1 pathway that controls *C. elegans* actomyosin contractility during embryonic development. SAX-3 is proposed to negatively regulate HUM-7, which functions as a GAP to attenuate RHO-1 activity and thus reduce actomyosin contractility through NMY-2, non-muscle myosin heavy chain 2. NMY-2 promotes acto-myosin contractility that is here proposed to be tightly regulated to permit cell migrations of ventral enclosure (Piekny et al., 2003).

The effects of Rho signaling on cell migration might occur through the activation of actomyosin contractility under the control of non-muscle myosin, NMY-2 in the epidermis. We crossed the *nmy-2::gfp* strain to *Plin-26::Lifeact::mCherry* to visualize the epidermis, and detected *nmy-2::gfp* puncta in the epidermal focal plane (Figure 5C), as previously shown by Wernike and colleagues (Wernike et al., 2016). During ventral enclosure, myosin puncta are enriched towards the leading edge of migrating cells, and at the midline as the cells meet(Fig. 5C). We examined how loss of *hum-7* affected *nmy-2::gfp* levels. Overall, all tissues of the embryo showed brighter *nmy-2::gfp* puncta in *hum-7* RNAi and mutant embryos. To measure effects on ventral enclosure, we measured the robust *nmy-2::gfp* epidermal signal as the pocket cells met at the midline. Plotting the maximum intensity showed an increase of 20% in *hum-7* mutants relative to controls (Fig. 5B).

We tested how two proposed upstream regulators of *hum-7* affected *nmy-2::gfp* levels during ventral pocket cell meeting. A null mutation in *sax-3*, *ky123*, led to a 20% decrease in the pocket cell signal while a null mutation in *vab-1*, *dx31*, led to 20% increase (Figure 5B,D). These results suggest both SAX-3 and VAB-1 regulate actomyosin contractility in the migrating epidermal cells. To test if *hum-7* is required for these changes we removed *hum-7* via RNAi in each strain. In the *vab-1(dx31); hum-7* RNAi strain the myosin levels are somewhere in the middle, closer to wild type, which suggests the effects of VAB-1 on HUM-7 may be indirect. In contrast, the levels of *nmy-2::gfp* in the *sax-3(ky123); hum-7(RNAi)* strain resembled the levels of *hum-7(RNAi)* (elevated) further supporting that SAX-3 acts through HUM-7.

## Discussion

### HUM-7, a new component in the RHO-1 pathway that appears to attenuate RHO signaling

The analysis of Rho GAPs has been complicated by their sheer number. Even in *C. elegans* which has fewer GAPs than mammals, there are 23 proteins with GAP domains to regulate the small number of small GTPases. Our analysis of one of these 23 GAP proteins has revealed exciting connections between axonal guidance receptors, the known players in RHO-1 signaling, and the movements of the epidermis during ventral enclosure. Our findings support a requirement for RHO-1 attenuation by the previously uncharacterized RhoGAP, HUM-7/Myosin 9. *hum-7* mutants have low penetrance ventral enclosure defects, coupled with highly penetrant increased actin dynamics, which suggests that increased actin dynamics in epidermal protrusions interferes with the normal processes that accompany cellular migration. It has been shown in single cell migration systems that down regulation of actomyosin contractility is required to allow persistent migration in one direction (Theisen et al., 2012). A similar event may be occurring in the migration of single axons, as we show here (Fig. 1) that loss of H M-7 alters neuronal migrations. One interpretation of our results is that in a sheet migration of the embryonic epidermis, increased actomyosin contractility interferes with persistent migration. Therefore increased actin dynamics, in the context of an embryo, reveals occasional faster migrations, but also failed migrations, likely due to lack of persistence in the direction of the migration direction.

### Axonal guidance receptors regulate the RHO-1 pathway

Our results suggest that HUM-7 plays an important role coordinating the response to external polarity cues by modulating RHO-1 activity to permit the correct levels and dynamics of actin required for proper cellular movements. HUM-7’s role in embryonic morphogenesis was discovered through a screen for enhancers of the partially penetrant morphogenesis defects caused by loss of an axonal guidance receptor, UNC-40/DCC (Fig.1). *hum-7* showed distinct genetic interactions with each of three axonal guidance pathway receptors.

Particularly intriguing was the rescue of embryonic lethality for *sax-3(ky123)* null animals, from 44% to 29% when *hum-7* is removed via RNAi or genetic mutations (Figure 1B, 2C). Since HUM-7 is proposed to regulate the RHO-1 pathway, we tested if the axonal guidance mutants affected embryonic non-muscle myosin, using the *nmy-2::gfp* CRISPR strain. The interaction was particularly interesting for SAX-3/ROBO. Loss of SAX-3 alters NMY-2::GFP levels, and this is epistatic to *hum-7*, suggesting *sax-3* and *hum-7* are in a pathway that regulates NMY-2. The connection between SAX-3 and HUM-7 is further supported by the fact that loss of *sax-3* results in elevated expression of overall GFP::HUM-7 levels measured using a CRISPR line (Fig. 3E). Our results support that SAX-3 is a positive regulator of RHO-1 signaling during the migrations of epidermal enclosure, and that SAX-3 acts, at least in part, through negative regulation of the RhoGAP HUM-7. Since loss of *hum-7* does not fully rescue loss of SAX-3, it is clear SAX-3 has other targets during ventral enclosure.

It is possible that VAB-1/EphrinB also regulates HUM-7, to regulate the RHO-1 pathway. Overall *vab-1* has opposite effects as *sax-3* in combination with RHO-1 pathway mutants. For example, loss of Rho Kinase/LET-502, which reduces RHO-1 signaling, results in embryonic lethality that can be rescued by loss of *hum-7* or *vab-1*, but not by loss of *sax-3*, suggesting VAB-1 has an overall negative effect on RHO-1 signaling (Fig. 4B). In support of this, loss of *vab-1* enhances loss of Myosin Phosphatase/MEL-11, just like loss of *hum-7*, and similarly increased levels of *nmy-2::gfp*. However, the effects of VAB-1 on the RHO-1 pathway may not be directly through HUM-7. Loss of *vab-1* did not significantly affect *gfp::hum-7* levels (Fig. 3D,E).

### Axonal guidance receptors regulate the RHO-1 pathway in epidermis

During ventral enclosure, *rho-1* and *nmy-2* are clearly needed in the underlying neuroblasts, as shown by Alisa Piekny and colleagues (Wernike et al., 2016). Analysis of the pattern of *nmy-2::gfp* in combination with transgenes only expressed in the epidermis confirms there are important *nmy-2::gfp* puncta that form during ventral enclosure (Figure 4C). Our findings that these epidermal *nmy-2::gfp* puncta are altered in embryos missing the axonal guidance receptors, SAX-3/ROBO and VAB-1/EphB, supports an important role for these proteins in guiding the epidermal migrations. While it is accepted that SAX-3 can rescue epidermal migrations when expressed only in the epidermis or the underlying neuroblasts (Ghenea et al., 2005) how VAB-1 influences epidermal migrations is more controversial, since only a subset of epidermal cells are proposed to express VAB-1 (Ikegami et al., 2012). In this context, it is intriguing that the strongest HUM-7 expression is not in the epidermis, or neuroblasts, but in the underlying muscle cells (Figure 3).

### HUM-7 and RHO-1 pathway affect actin dynamics

SAX-3/ROBO regulation of HUM-7/Myo9 to regulate Rho is conserved from human cells (lung cancer) to *C. elegans*. The homologs of HUM-7, Myo9A and Myo9B, are proposed to regulate cellular behaviors as GAPs that act on the Rho GTPase. Many of the proposed phenotypes can be explained as resulting from excess Rho signaling. What is not as clear, is what are the downstream effects of this excess Rho signaling, and what signals regulate the behavior of this unusual RhoGAP with a processive myosin motor. We used a complex tissue migration, ventral enclosure, to measure several aspects of Rho signaling to ask how it can be perturb the dynamics of the tissue migration. As the first analysis of Myosin 9 function in *C. elegans*, we have uncovered many features of morphogenesis that depend on Myo9, and have begun to place them in signaling pathways. We describe that axonal guidance in *C. elegans* depends on HUM-7/Myosin9, which appears to receive signals through SAX-3/ROBO, and possibly VAB-1/EphrinB. These interactions are conserved in embryos, where they regulate epidermal cell migrations. The signals, originating from the axonal guidance receptors, alter non-muscle myosin expression in the migrating epidermal cells. The pathway we uncover here, from SAX-3/ROBO to HUM-7/Myo9, to modulate RHO-1/Rho, has been proposed to function in human lung cancer tissue culture cells. The altered timing of the migrations in *hum-7* mutant embryos suggests one consequence of misregulated Rho in the absence of this RhoGAP is overactive Rho that leads to excess protrusions at the leading edge, without a matching increase in retractions. This change may create various problems. It is possible the protrusions are too stable, due to decreased turnover of Rho-GTPase. While this may lead to occasional faster migrations, it may also make the cells less responsive to their environment. For migrating epidermal cells, this could reduce detection of tension in the underlying neuroblasts that are thought to help guide the migrations. The defects in *hum-7* mutants, therefore, may combine failure to properly transmit signals within the epidermal cells, and also, failure to detect signals from cells in neighboring tissues. Tissue specific rescue experiments of *hum-7* to resolve this would be challenging, since the overall lethality is so low. However, future experiments will need to address the tissue specificity of the signals, and of HUM-7 function in receiving these signals, to address how the epidermis and neuroblasts cooperate during this complex migration.

## Materials and Methods

### Strains

The following strains were used in this study: MT324 *unc-40(n324)*, CX3198 *sax-3(ky123)*, CZ337 *vab-1(dx31)*, FT48 *xnls16[dlg-1::gfp]*, VC2436 *hum-7(ok3054)*, OX615 *hum-7(ok3054); dlg-1::gfp*, WM43 *gex-3(zu196)/DnT 1*, MT5013 *ced-10(n1993)*, MT9958 *ced-10(n3246)*, JH2754 *ect-2(ax751)*, OX646 *hum-7(ok3054); ect-2(ax751)*, JJ1473 *zuls45[nmy-2::gfp]*, OX630 *hum-7(ok3054); nmy-2::gfp*, ML1154 *mcls51[plin26::vab-10 ABD::gfp]*, OX595 *hum-7(ok3054); plin26::vab-10 ABD::gfp*, SK4005 *zdls5[mec-4::gfp]*, OX350 *unc-40(n324); mec-4::gfp*, OX213 *sax-3(ky123); mec-4::gfp*, IC136 *vab-1(dx31); mec-4::gfp*, HR1157 *let-502(sb118ts)*, OX644 *hum-7(ok3054) let-502(sb118ts)*, OX635 *hmp-1(fe4); hum-7(ok3054)*, OX681 *gfp::hum-7*, OX714 *hum-7(pj62)*, OX715 *hum-7(pj63)*, ML773 *rga-2(hd102)/hln1 [unc-54(h1040)]*, OX645 *hum-7(ok3054) rga-2(hd102)/hln1 [unc-54(h1040)]*, HR483 *mel-11(sb56) unc-4(e120)/mnC1[dpy-10(e128) unc-52(e444)]*.

### ln vitro binding assay

This assay was based on (Neukomm et al., 2011). GTP-loaded and GDP-loaded GTPase constructs for CDC-42, CED-10 and RHO-1 were gifts from the Hengartner lab. The HUM-7 GAP domain construct (aa1540-1727) was cloned into the pGEX-4T2 GST vector. All the GTPase constructs were His-tagged and purified using His-Bind resin (Novagen). The GST-tagged HUM-7 construct was purified using Glutathione Sepharose 4B beads (GE Healthcare) based on manufacturers instructions. For the pull-down assay, 30μg of purified GST-tagged proteins were incubated with 10μg His-tagged proteins at 4°C for two hours. Pull-downs were performed with GST beads, proteins were separated on 12% acrylamide gels, and blots were probed with antibody to His (Millipore).

### RNA interference

All RNAi used in this study was administered by the feeding protocol as in (Bernadskaya et al., 2012). RNAi were constructed by cloning cDNA’s of the genes into the L4440 vector and transforming them into HT115 competent cells. Small overnight bacterial cultures were diluted 1:250 and grown until the OD600 was close to 1. The culture was pelleted and resuspended in LB media containing 100 pg/ml Ampicillin and 1mM IPTG.

### Embryonic Lethality counts and imaging

Synchronized L1 worms were plated on AMP/IPTG plates containing the appropriate RNAi bacteria. Plates were cultured at 23°C for three days. Temperature sensitive mutants like *ect-2(ax751)* were cultured at three different temperatures, 15°C, 20°C and 25°C. After the three-day incubation, embryos were mounted on 3% agarose pads and lethality was counted.

### Neuronal Migration Analysis

For *mec-4::gfp* neuronal analysis, synchronized L1 worms were plated on AMP/IPTG plates containing the appropriate RNAi bacteria and cultured for 2-3 days at 20°C. L4440 empty vector was used as the control RNAi. The effectiveness of *hum-7* RNAi was monitored by counting the percent embryonic lethality. After incubation, L4 worms were mounted unto 3% agarose pads with 10mM levamisole to prevent frequent movement and scored for phenotypes within 15 minutes of mounting. The AVM mechanosensory neuron was checked for ventral migration defects. The table in Fig. 2B is based on more than 200 worms per genotype.

### RNAi-feeding Screen

To generate liquid cultures of the RNAi genes, 150μl of LB broth containing the appropriate antibiotic was pipetted into 96-well plates. Using a 96-well pin, frozen glycerol stocks of RNAi genes were transferred to the liquid media. The culture was inoculated overnight at 37°C with shaking. Cultures were spotted on 24-well NGM plates containing 2% lactose and 25μg/ml carbenicillin and grown overnight at room temperature. Approximately 20 L1 worms with the *unc-40(n324)* genotype were seeded unto the 24-well plates containing the RNAi bacteria. Worms were grown at 23°C for about three days and then their progeny screened for embryonic lethality. Clones that produced greater than 15% lethality in combination with *unc-40(n324)* were selected for secondary screening. We were particularly interested in selecting clones that had good differentiation and phenotypes that resemble Gex.

### Live lmaging

Some late stage *hum-7* embryos (<2%) showed a subtle phenotype in which they developed correctly but flattened when mounted on pads for imaging. This flat appearance may be related to the fact that *hum-7* embryos behave osmotically sensitive, and have to be mounted in Egg Salts instead of water to prevent arrest during live imaging studies. Thus all embryos shown in this study were mounted in Egg Salts.

To measure actin levels and dynamics, we performed live imaging of the *plin-26;vab-10ABD::GFP* transgene (Gally et al., 2009). Embryos at the 2 to 4 cell stage were dissected from adult hermaphrodites and mounted onto 3% agarose pads. Embryos were then incubated at 23°C for 240 minutes. Following the 240-minute incubation, embryos were imaged at 2-minute intervals for 120 minutes. Imaging was done on a Laser Spinning Disk Confocal Microscope with a Yokogawa scanhead, on a Zeiss AxioImager Z1m Microscope using the Plan-Apo 63X/1.4NA or Plan-Apo 40X/1.3NA oil lenses. Images were captured on a Photometrics Evolve 512 EMCCD Camera using MetaMorph software, and analyzed using ImageJ.

### Statistical Analysis

All graphs show the mean of the data and the Standard Error of the Mean (SEM). For grouped data, statistical significance was established by performing a two-way Analysis of Variance (ANOVA) followed by the Bonferroni multiple comparison posttest. For ungrouped data a one-way ANOVA was performed followed by the Tukey post-test. Asterisks (*) denote p values <0.05. All statistical analysis was performed using GraphPad Prism.

## Acknowledgments

We would like to thank the NCRR-funded *Caenorhabditis* Genetics center, Jeremy Nance, and Paul Mains for strains, and Lucas Neukomm for GTPase constructs. We thank Rutgers undergraduate Vipin Palukuri for technical assistance. Thanks to Chris Rongo, Monica Driscoll and Andy Singson for helpful comments on the manuscript and Alisa Piekny for helpful discussions.

## Supporting Information

**S1 Movie. F-actin dynamics and levels in *wild type* vs. *hum-7(ok3054)* embryos**. The epidermal-specific transgene, *plin-26:: vab-10 ABD::gfp*, was used to image the F-actin dynamics in the migrating epidermal cells during embryonic enclosure. Embryos were staged from the 2-cell stage and imaged at 23°C under identical conditions. Ventral is up. In this movie the entire embryos are shown. The center *hum-7(ok3054)* embryo encloses faster than wild type while the *hum-7(ok3054)* embryo on the right arrests partway through enclosure. In the right panel the bright signal at the lower left corner is from another embryo.

**S2 Movie. F-actin dynamics at the epidermal leading edge in a *wild type* vs. *hum-7(ok3054)* embryos**. A subsection of the same embryos from Movie S1 is shown to focus on the dynamics at the leading edge of the migrating ventral cells.

## References

Abouhamed, M., Grobe, K., Leefa Chong San, I.V., Thelen, S., Honnert, U., Balda, M.S., Matter, K., Bähler, M., 2009. Myosin IXa Regulates Epithelial Differentiation and Its Deficiency Results in Hydrocephalus. Molecular Biology of the Cell 20, 5074–5085.

Bähler, M., Rhoads, A., 2002. Calmodulin signaling via the IQ motif. FEBS Letters513, 107–113.

Bernadskaya, Y.Y., Wallace, A., Nguyen, J., Mohler, W.A., Soto, M.C., 2012. UNC-40/DCC, SAX-3/Robo, and VAB-1/Eph Polarize F-Actin during Embryonic Morphogenesis by Regulating the WAVE/SCAR Actin Nucleation Complex. PLoS Genetics 8, e1002863.

Chandhoke, S.K., Mooseker, M.S., 2012. A role for myosin IXb, a motor-RhoGAP chimera, in epithelial wound healing and tight junction regulation. Molecular Biology of the Cell23, 2468–2480.

Chisholm, A.D., Hardin, J., 2005. Epidermal morphogenesis.

Clay, M.R., Halloran, M.C., 2013. Rho activation is apically restricted by Arhgap1 in neural crest cells and drives epithelial-to-mesenchymal transition. Development (Cambridge, England)140, 3198–3209.

Diogon, M., Wissler, F., Quintin, S., Nagamatsu, Y., Sookhareea, S., Landmann, F., Hutter, H., Vitale, N., Labouesse, M., 2007. The RhoGAP RGA-2 and LET-502/ROCK achieve a balance of actomyosin-dependent forces in C. elegans epidermis to control morphogenesis. Development134, 2469–2479.

Fotopoulos, N., Wernike, D., Chen, Y., Makil, N., Marte, A., Piekny, A., 2013. Caenorhabditis elegans anillin (ani-1) regulates neuroblast cytokinesis and epidermal morphogenesis during embryonic development. Developmental biology383, 61–74.

Gally, C., Wissler, F., Zahreddine, H., Quintin, S., Landmann, F., Labouesse, M., 2009. Myosin II regulation during C. elegans embryonic elongation: LET-502/ROCK, MRCK-1 and PAK-1, three kinases with different roles. Development136, 3109–3119.

Ghenea, S., Boudreau, J.R., Lague, N.P., Chin-Sang, I.D., 2005. The VAB-1 Eph receptor tyrosine kinase and SAX-3/Robo neuronal receptors function together during <em>C. elegans</em> embryonic morphogenesis. Development132, 3679–3690.

Hanley, P.J., Xu, Y., Kronlage, M., Grobe, K., Schön, P., Song, J., Sorokin, L., Schwab, A., Bähler, M., 2010. Motorized RhoGAP myosin IXb (Myo9b) controls cell shape and motility. Proceedings of the National Academy of Sciences107, 12145–12150.

Havrylenkoxs, S., Noguera, P., Abou-Ghali, M., Manzi, J., Faqir, F., Lamora, A., Guérin, C., Blanchoin, L., Plastino, J., 2015. WAVE binds Ena/VASP for enhanced Arp2/3 complex-based actin assembly. Molecular Biology of the Cell26, 55–65.

Hegan, P.S., Chandhoke, S.K., Barone, C., Egan, M., Bähler, M., Mooseker, M.S., 2016. Mice lacking myosin IXb, an inflammatory bowel disease susceptibility gene, have impaired intestinal barrier function and superficial ulceration in the ileum. Cytoskeleton73, 163–179.

Ikegami, R., Simokat, K., Zheng, H., Brown, L., Garriga, G., Hardin, J., Culotti, J., 2012. Semaphorin and Eph Receptor Signaling Guide a Series of Cell Movements for Ventral Enclosure in C. elegans. Current Biology22, 1–11.

Inoue, A., Saito, J., Ikebe, R., Ikebe, M., 2002. Myosin IXb is a single-headed minus-end-directed processive motor. Nature Cell Biology 4, 302.

Kong, R., Yi, F., Wen, P., Liu, J., Chen, X., Ren, J., Li, X., Shang, Y., Nie, Y., Wu, K., Fan, D., Zhu, L., Feng, W., Wu, J.Y., 2015. Myo9b is a key player in SLIT/ROBO-mediated lung tumor suppression. The Journal of Clinical Investigation125, 4407–4420.

Li, P., Yang, X.-K., Wang, X., Zhao, M.-Q., Zhang, C., Tao, S.-S., Zhao, W., Huang, Q., Li, L.-J., Pan, H.-F., Ye, D.-Q., 2016. A meta-analysis of the relationship between MYO9B gene polymorphisms and susceptibility to Crohn’s disease and ulcerative colitis. Human Immunology77, 990–996.

Liao, W., Elfrink, K., Bähler, M., 2010. Head of Myosin IX Binds Calmodulin and Moves Processively toward the Plus-end of Actin Filaments. The Journal of Biological Chemistry285, 24933–24942.

Makowska, Katarzyna A., Hughes Ruth E., White, Kathryn J., Wells, Claire M., Peckham, M., 2015. Specific Myosins Control Actin Organization, Cell Morphology, and Migration in Prostate Cancer Cells. Cell Reports13, 2118–2125.

Marston, Daniel J., Higgins, Christopher D., Peters, Kimberly A., Cupp, Timothy D., Dickinson, Daniel J., Pani, Ariel M., Moore, Regan P., Cox, Amanda H., Kiehart, Daniel P., Goldstein, B., 2016. MRCK-1 Drives Apical Constriction in C. elegans by Linking Developmental Patterning to Force Generation. Current Biology26, 2079–2089.

Martin, E., Harel, S., Nkengfac, B., Hamiche, K., Neault, M., Jenna, S., 2014. pix-1 Controls Early Elongation in Parallel with mel-11 and let-502 in Caenorhabditis elegans. PLoS ONE 9, e94684.

Miller, D.M. III, Ortiz, I., Berliner, G.C., Epstein, H.F., 1983. Differential localization of two myosins within nematode thick filaments. Cell34, 477–490.

Neukomm, L.J., Frei, A.P., Cabello, J., Kinchen, J.M., Zaidel-Bar, R., Ma, Z., Haney, L.B., Hardin, J., Ravichandran, K.S., Moreno, S., Hengartner, M.O., 2011. Getting rid of damaged goods: Loss of RhoGAP SRGP-1 promotes the clearance of dead and injured cells in C. elegans. Nature Cell Biology 13, 10.1038/ncb2138.

Neukomm, L.J., Zeng, S., Frei, A.P., Huegli, P.A., Hengartner, M.O., 2014. Small GTPase CDC-42 promotes apoptotic cell corpse clearance in response to PAT-2 and CED-1 in C. elegans. Cell Death and Differentiation21, 845–853.

O’Connor, E., Töpf, A., Müller, J.S., Cox, D., Evangelista, T., Colomer, J., Abicht, A., Senderek, J., Hasselmann, O., Yaramis, A., Laval, S.H., Lochmüller, H., 2016. Identification of mutations in the MYO9A gene in patients with congenital myasthenic syndrome. Brain139, 2143–2153.

Omelchenko, T., Hall, A., 2012. Myosin-IXA Regulates Collective Epithelial Cell Migration by Targeting RhoGAP Activity to Cell-Cell Junctions. Current Biology22, 278–288.

Ouellette, M.-H., Martin, E., Lacoste-Caron, G., Hamiche, K., Jenna, S., 2016. Spatial control of active CDC-42 during collective migration of hypodermal cells in Caenorhabditis elegans. Journal of Molecular Cell Biology8, 313–327.

Patel, F.B., Bernadskaya, Y.Y., Chen, E., Jobanputra, A., Pooladi, Z., Freeman, K.L., Gally, C., Mohler, W.A., Soto, M.C., 2008. The WAVE/SCAR complex promotes polarized cell movements and actin enrichment in epithelia during C. elegans embryogenesis. Developmental biology324, 297–309.

Piekny, A., Mains, P., 2003. Squeezing an egg into a worm: C. elegans embryonic morphogenesis. The Scientific World Journal3, 1370–1381.

Piekny, A.J., Johnson, J.-L.F., Cham, G.D., Mains, P.E., 2003. The Caenorhabditis elegans nonmuscle myosin genes nmy-1 and nmy-2 function as redundant components of the let-502/Rho-binding kinase and mel-11/myosin phosphatase pathway during embryonic morphogenesis. Development130, 5695–5704.

Piekny, A.J., Wissmann, A., Mains, P.E., 2000. Embryonic morphogenesis in Caenorhabditis elegans integrates the activity of LET-502 Rho-binding kinase, MEL-11 myosin phosphatase, DAF-2 insulin receptor and FEM-2 PP2c phosphatase. Genetics156, 1671–1689.

Prager, M., Durmus, T., Büttner, J., Molnar, T., de Jong, D.J., Drenth, J.P.H., Baumgart, D.C., Sturm, A., Farkas, K., Witt, H., Büning, C., 2014. Myosin IXb variants and their pivotal role in maintaining the intestinal barrier: A study in Crohn’s disease. Scandinavian Journal of Gastroenterology49, 1191–1200.

Quintin, S., Gally, C., Labouesse, M., 2008. Epithelial morphogenesis in embryos: asymmetries, motors and brakes. Trends in Genetics24, 221–230.

Sieburth, D., Ch’ng, Q., Dybbs, M., Tavazoie, M., Kennedy, S., Wang, D., Dupuy, D., Rual, J.F., Hill, D.E., Vidal, M., Ruvkun, G., Kaplan, J.M., 2005. Systematic analysis of genes required for synapse structure and function. Nature436, 510–517.

Soto, M.C., Qadota, H., Kasuya, K., Inoue, M., Tsuboi, D., Mello, C.C., Kaibuchi, K., 2002. The GEX-2 and GEX-3 proteins are required for tissue morphogenesis and cell migrations in C. elegans. Genes & Development16, 620–632.

Theisen, U., Straube, E., Straube, A., 2012. Directional Persistence of Migrating Cells Requires Kif1C-Mediated Stabilization of Trailing Adhesions. Developmental cell23, 1153–1166.

Thelen, S., Abouhamed, M., Ciarimboli, G., Edemir, B., Bähler, M., 2015. Rho GAP myosin IXa is a regulator of kidney tubule function. American Journal of Physiology-Renal Physiology 309, F501–F513.

Viveiros, R., Hutter, H., Moerman, D.G., 2011. Membrane extensions are associated with proper anterior migration of muscle cells during Caenorhabditis elegans embryogenesis. Developmental Biology358, 189–200.

Walck-Shannon, E., Lucas, B., Chin-Sang, I., Reiner, D., Kumfer, K., Cochran, H., Bothfeld, W., Hardin, J., 2016. CDC-42 Orients Cell Migration during Epithelial Intercalation in the Caenorhabditis elegans Epidermis. PLoS Genetics 12, e1006415.

Wang, M.-J., Xu, X.-L., Yao, G.-L., Yu, Q., Zhu, C.-F., Kong, Z.-J., Zhao, H., Tang, L.-M., Qin, X.-H., 2016. MYO9B gene polymorphisms are associated with the risk of inflammatory bowel diseases. Oncotarget7, 58862–58875.

Wernike, D., Chen, Y., Mastronardi, K., Makil, N., Piekny, A., 2016. Mechanical forces drive neuroblast morphogenesis and are required for epidermal closure. Developmental Biology412, 261–277.

Wissmann, A., Ingles, J., McGhee, J.D., Mains, P.E., 1997. Caenorhabditis elegans LET-502 is related to Rho-binding kinases and human myotonic dystrophy kinase and interacts genetically with a homolog of the regulatory subunit of smooth muscle myosin phosphatase to affect cell shape. Genes & Development11, 409–422.

Yu, T.W., Hao, J.C., Lim, W., Tessier-Lavigne, M., Bargmann, C.I., 2002. Shared receptors in axon guidance: SAX-3/Robo signals via UNC-34/Enabled and a Netrin-independent UNC-40/DCC function. Nat Neurosci5, 1147–1154.

Zaidel-Bar, R., Joyce, M.J., Lynch, A.M., Witte, K., Audhya, A., Hardin, J., 2010. The F-BAR domain of SRGP-1 facilitates cell-cell adhesion during <em>C. elegans</em> morphogenesis. The Journal of Cell Biology191, 761–769.

Zhang, H., Gally, C., Labouesse, M., 2010. Tissue morphogenesis: how multiple cells cooperate to generate a tissue. Current Opinion in Cell Biology22, 575–582.

Zilberman, Y., Abrams, J., Anderson, D.C., Nance, J., 2017. Cdc42 regulates junctional actin but not cell polarization in the <em>Caenorhabditis elegans</em> epidermis. The Journal of Cell Biology.

Zonies, S., Motegi, F., Hao, Y., Seydoux, G., 2010. Symmetry breaking and polarization of the C. elegans zygote by the polarity protein PAR-2. Development (Cambridge, England)137, 1669–1677.

